# Proteomic Analysis Reveals Trilaciclib-Induced Senescence

**DOI:** 10.1101/2024.03.12.584620

**Authors:** Marina Hermosilla-Trespaderne, Mark Xinchen Hu-Yang, Abeer Dannoura, Andrew M. Frey, Amy L. George, Matthias Trost, José Luis Marín-Rubio

## Abstract

Trilaciclib, a CDK4/6 inhibitor, was approved as a myeloprotective agent for protecting bone marrow from chemotherapy-induced damage in extensive-stage small cell lung cancer (ES-SCLC). This is achieved through the induction of a temporary halt in the cell cycle of bone marrow cells. While it has been studied in various cancer types, its potential in haematological cancers remains unexplored. This research aimed to investigate the efficacy of trilaciclib in haematological cancers. Utilizing mass spectrometry-based proteomics, we examined the alterations induced by trilaciclib in the chronic myeloid leukaemia (CML) cell line, K562. Interestingly, trilaciclib promoted senescence in these cells rather than cell death, as observed in acute myeloid leukaemia (AML), acute lymphoblastic leukaemia (ALL), and myeloma cells. In K562 cells, trilaciclib hindered cell cycle progression and proliferation by stabilising CDK4/6 and downregulating cell cycle-related proteins, along with the concomitant activation of autophagy pathways. Additionally, trilaciclib-induced senescence was also observed in the non-small cell lung carcinoma cell line (NSCLC), A549. These findings highlight trilaciclib’s potential as a therapeutic option for haematological cancers and underscore the need to carefully balance senescence induction and autophagy modulation in CML treatment, as well as in NSCLC.

**ABSTRACT GRAPHIC:** 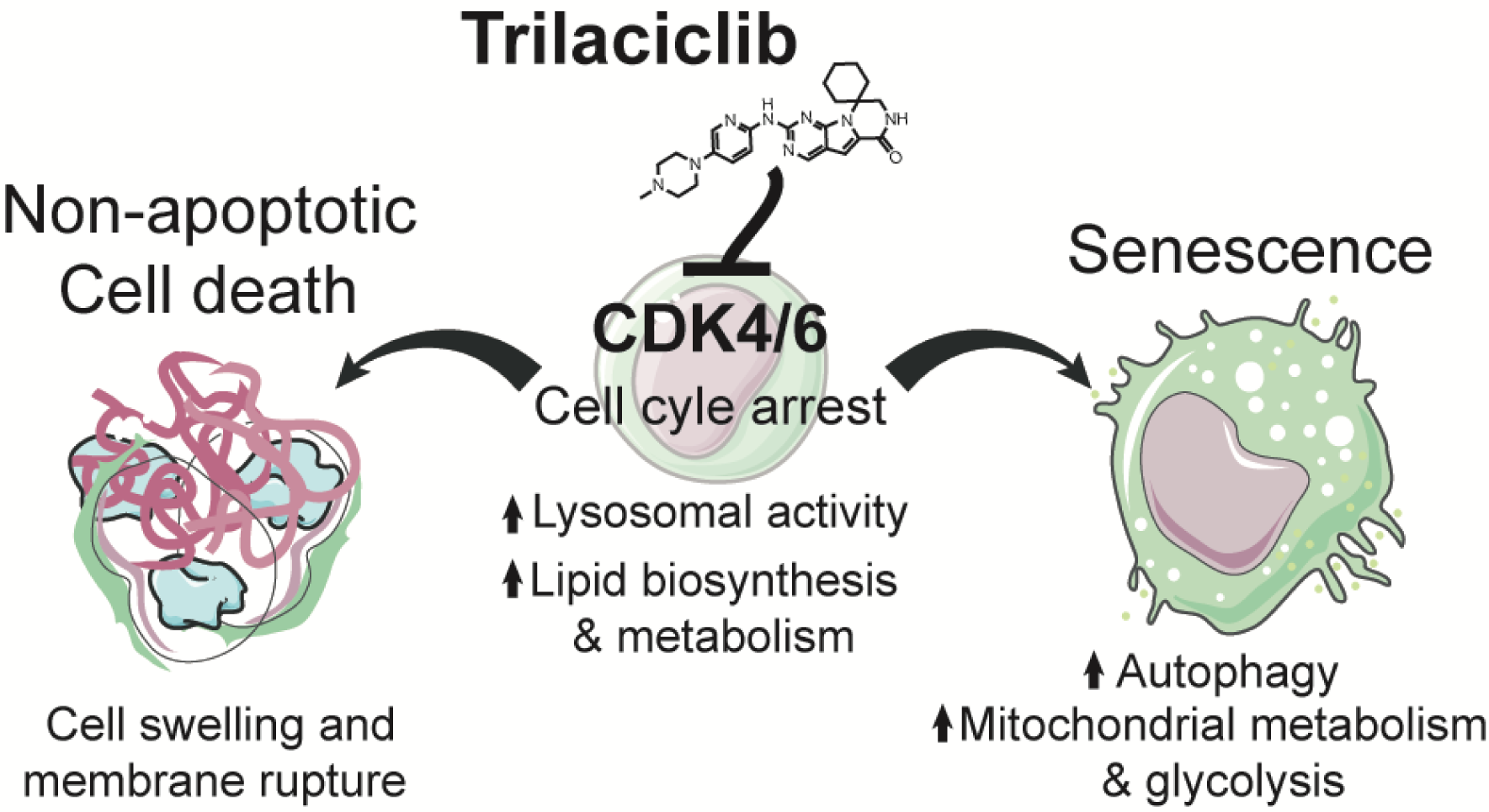

## INTRODUCTION

Trilaciclib was approved by the FDA in 2021 as a myeloprotective agent for adult patients with extensive-stage small cell lung cancer (ES-SCLC) (1,2). Trilaciclib is given prior to the administration of chemotherapy in order to prepare and protect the bone marrow from the potential damage caused by the chemotherapy treatment. Trilaciclib blocks cyclin-dependent kinase (CDK) 4/6, CDK4/6, which plays a crucial role in cell cycle progression, particularly in the transition from the G1 phase to the S phase. Clinical trials have evaluated the use of trilaciclib in various cancer types, including small cell lung cancer (SCLC) (3), advanced and metastatic triple-negative breast cancer (4,5), advanced and metastatic bladder cancer (NCT04887831), metastatic colorectal cancer (NCT04607668), advanced and metastatic non-small cell lung cancer (NSCLC) (NCT05900921 and NCT04863248, respectively). Recent analysis indicates that ES-SCLC patients treated with trilaciclib demonstrated lower performance status as assessed by the Eastern Cooperative Oncology Group (ECOG) score (6). Nevertheless, this compound has only been characterised to its target disease and has awakened the desire to put it to the test in other types of cancer such as haematological cancers or in tumoral cells of NSCLC.

Chronic myeloid leukaemia (CML) is mainly characterised by the presence of the Philadelphia chromosome, which produces the BCR-ABL oncoprotein (7). This oncoprotein is a constitutively active tyrosine kinase that promotes cell proliferation and survival through various signalling pathways (8). Targeted therapy with tyrosine kinase inhibitors (TKIs) has transformed CML into a manageable disease. To date, six TKIs have been approved for targeting the BCR-ABL oncoprotein: imatinib (first generation), dasatinib, nilotinib, and bosutinib (second generation), ponatinib and asciminib (third generation). While TKIs effectively control CML and prolong patient survival, they may not completely eradicate leukemic cells (9), leading to treatment failure and relapse. Furthermore, CML treatments can also cause chemotherapy-induced myelosuppression which is characterised by a decrease in blood cell counts (10,11).

The induction of senescence by leukemogenic fusion proteins (12–14), chemotherapy (12), or TKIs (15) in CML involves both positive and negative effects (16). Senescent cells exhibit cell cycle arrest and can suppress the growth and proliferation of leukemic cells. They can also stimulate immune responses, leading to the clearance of leukemic cells by the immune system. However, the failure to clear senescent cells can result in chronic inflammation, creating a tumour-promoting microenvironment. While senescence can contribute to disease control, the secretion of inflammatory molecules by senescent cells can have long-term side effects (17). In addition, autophagy can modulate cellular senescence which eliminates dysfunctional components within lysosomes during periods of cellular stress (18). Therefore, a careful balance is necessary in managing senescence induction in cancer treatment. Furthermore, understanding the senescence signature and the specific proteins that are downregulated or upregulated in senescent cells can provide insights into the molecular mechanisms underlying cellular ageing and senescence-associated diseases. Here, we showed the potential of trilaciclib inducing cell death in haematological cancers, particularly in acute myeloid leukaemia (AML), acute lymphoblastic leukaemia (ALL), myeloma, but not in the chronic myeloid leukaemia (CML) cell line, K562. We used proteomics-based mass spectrometry to characterise trilaciclib, observing that it is inducing senescence and autophagy in K562 cells. Additionally, the induction of senescence in the non-small cell lung cancer (NSCLC) cell line, A549, by trilaciclib suggests that this drug may exert its effects through alternative mechanisms in different cancer types. Overall, these findings suggest that trilaciclib holds promise as a potential therapeutic option for various cancer types, including haematological cancers like AML, ALL, and myeloma, and possibly even NSCLC.

## METHODS

### Cell lines

JURKAT (TIB-152), K562 (CCL-234), MOLT-4 (CRL-1582), NCI-H929 (CRL-9068), U937 (CRL-1593.2), and A549 (CCL-185) cell lines were purchased from the American Type Culture Collection (ATCC). ATCC routinely performs cell line authentication, using short tandem repeat profiling as a procedure. JURKAT, K562, NCI-H929, U937, and A549 cell lines were cultured in Roswell Park Memorial Institute medium (RPMI, Gibco) supplemented with 10 % foetal bovine serum (FBS) and 2 mM L-glutamine. Cell lines were grown at 37 °C in a humidified incubator containing 5 % CO_2_. Experiments with cell lines were performed within a period not exceeding three months after resuscitation and lines were regularly tested for mycoplasma.

### Drugs

Trilaciclib and palbociclib were purchased from SelleckChem. Oligomycin A (OM), carbonyl cyanide 4 (trifluoromethoxy) phenylhydrazone (FCCP), antimycin A (AA), rotenone (Rot), and 2-deoxyglucose (2-DG) were purchased from Sigma-Aldrich.

### *In silico* analysis of published datasets

Human gene mRNA expressions data were obtained from the Human Protein Atlas (version 23.0, available at https://www.proteinatlas.org) (19). Sensitivity of cancer cell lines to trilaciclib, imatinib, asciminib, and palbociclib data were acquired from the Cancer Dependency Map (DepMap, BRD:BRD-K00003412-300-01-9, BRD:BRD-K92723993-001-17-4, BRD:BRD-K00003456-001-01-9, and BRD:BRD-K51313569-003-03-3, PRISM Repurposing Public 23Q2; accessible at https://depmap.org/portal/) and the CRISPR analysis was conducted using the CRISPR-cas9 screening data from the Cancer Dependency Map generated at the Broad Institute (DepMap Public 21Q3+Score, Chronos; accessible at https://depmap.org/portal/) (20).

### Cell viability assays and IC50 trilaciclib concentration

The cells were treated with inhibitors for 72 hours, and the cell viability was measured by flow cytometry using propidium iodide solution (PI) (Millipore). The BD Symphony A5 cytometer was used in high-throughput mode to analyse 96-well plates using an excitation 560 nm and an emission of 610/20 nm. FlowJo Software (v10.8.1) was used to analyse the data. IC50 curves were calculated using different concentrations of trilaciclib (10, 7.5, 5, 2.5, 1, 0.8, 0.6, 0.4, 0.2, 0.1, 0.08, 0.06, 0.04, 0.02, and 0.01 μM) for 72 hours.

### Beta-galactosidase (SA-b-gal) activity assay

Treated cells were transferred to a poly-L-lysine treated 24-well plate and centrifuged at 1,000 rpm (112 ×g) for 5 min to allow the cells to adhere to the bottom of the wells. The SA-b-gal staining kit (#9860, Cell Signaling) was performed according to the manufacturer’s protocol. In short, the cells were incubated with the SA-b-gal solution at 37 °C for 72 hours. Following incubation, the cells were washed twice with PBS, covered with 70 % glycerol, and stored at 4 °C. Nikon Tie microscope was used for photographic material acquisition at a 10× magnification. The percentages of positive cells were calculated from a total of one hundred cells regardless of their intensity, across four biological replicates. Imaging analysis was conducted using ImageJ (Fiji 2.0.0).

### LysoTracker

The cells were treated with inhibitors for 24 hours, 3 and 5 days and stained with 50 nM LysoTracker Red DND-99 for 2 hours. Four independent experiments were analysed. All analytic cytometry procedures were performed on a BD Symphony A5 cytometer (Becton Dickinson). The analysis was performed using FlowJo (v10.8.1).

### Proliferation assay

The cells were treated with inhibitors for 12 or 7 days, and cell counting was assessed using dye exclusion with Trypan Blue Solution, 0.4 % (Thermo Fisher). Three independent experiments were analysed.

### Cell death assay

The cells were stained using the PE Annexin V Apoptosis Detection Kit (BD Pharmingen) according to the manufacturer’s protocol. The BD Symphony A5 cytometer was used in high-throughput mode to analyse 96-well plates using compensation between excitation 561 nm and an emission of 710/50 nm, and 561 nm and an emission 586/15 nm.

### Cell cycle analysis

Cell cycle profiling using PI/RNase Staining Buffer (BD Biosciences) was performed according to the manufacturer’s instructions. Phase distribution of the cell cycle was examined after 72 hours of treatments. All analytic cytometry procedures were performed on a BD Symphony A5 cytometer (Becton–Dickinson) in high-throughput mode. The cell cycle phase analysis was performed using FlowJo (v10.8.1).

### Seahorse metabolic analysis

The oxygen consumption rate (OCR) and the extracellular acidification rate (ECAR) were measured using Seahorse XFp Analyzer (Agilent Technologies). Treated cells were transferred to a poly-L-lysine treated 96-well plate and centrifuged at 1,000 rpm (112 ×g) for 5 minutes to allow the cells to adhere to the bottom of the wells. Then, the cell culture plate was placed into an incubator at 37 °C and without CO_2_. Three measurements were analysing in baseline conditions and after four sequentially injections of 15 µM oligomycin A (OM), 1.5 µM carbonyl cyanide 4 (trifluoromethoxy) phenylhydrazone (FCCP), 2.5 µM antimycin A (AA) with 1.25 µM rotenone (Rot), and finally 50 mM 2-deoxyglucose (2-DG) final concentration. ECAR and OCR were analysed using the Wave 2.6.3 software and the parameters were calculated as in previous publications (21).

### Cellular Thermal Shift Assay

Intact-cell experiments were performed on K562 cells as described previously, with some slight alterations (22). Briefly, 30 million K562 cells were treated with 1 μM trilaciclib or vehicle (0.01 % DMSO) in complete media at 37 °C for 1 hour. Cells were collected and washed twice with PBS containing 1 μM trilaciclib or vehicle. Washed cells were suspended in PBS supplemented with treatment or vehicle plus 0.4 % NP-40, cOmplete Protease Inhibitor Cocktail (Sigma-Aldrich) and phosphatase inhibitor cocktail (1.2 mM sodium molybdate, 1 mM sodium orthovanadate, 4 mM sodium tartrate dihydrate, and 5 mM glycerophosphate) and then separated into eight fractions for thermal profiling. Fractions were heated at 40, 44, 48, 52, 56, 60, 64, and 68 °C for 3 minutes, incubated for 3 minutes at room temperature, and snap-frozen at −80 °C. Samples were lysed with four freeze–thaw cycles using dry ice and a thermo block at 35 °C. Cell lysates were centrifuged at 15,000×g for 2 hours at 4 °C to separate protein aggregates from soluble proteins. Supernatants were collected for western blot.

### Protein lysis

Cells were lysed using lysis buffer (5 % SDS, 50 mM triethylammonium bicarbonate (TEAB) pH 7.55 supplemented with protease inhibitors (Complete, EDTA free protease inhibitor cocktail), phosphatase (1.15 mM sodium molybdate, 4 mM sodium tartrate dihydrate, 1 mM freshly prepared sodium orthovanadate, 5 mM glycerophosphate), and nucleases (Pearson)) and sonicated for 30 seconds.

### Western blot

Ten micrograms of proteins from each sample were separated on 10 % or 12 % sodium dodecyl-sulphate polyacrylamide gel electrophoresis gel before being transferred to polyvinylidene difluoride membranes. Ponceau red staining was performed after transference. After that, the membrane was blocked with 5 % w/v BSA in 0.1 % Tween-20 TBS 1X for 1 hour at room temperature. Then, it was probed with phospho-Rb (Ser 807/811) (#8516, Cell Signaling), p21 (#2947, Cell Signaling), CDK4 (sc-23896, Santa Cruz), CDK6 (#13331, Cell Signaling), α-Tubulin (T9026) (Sigma-Aldrich) antibodies overnight at 4 °C. Afterwards, membranes were incubated for 1 hour at room temperature with anti-rabbit IgG (#7074, Cell Signalling) or anti-mouse IgG (#7076) antibody. Pierce ECL Western Blotting Substrate (Thermo Scientific) was used for detection and Amersham GE Imager (GE Healthcare) for imaging the membranes through chemiluminescence.

### Sample preparation for LC-MS/MS

Samples were prepared for mass spectrometry using S-Trap micro columns (Protifi) according to the manufacturer recommended protocol. Proteins were reduced using 20 mM tris((2-carboxyethyl) phosphine) for 30 minutes at 47 °C with gentle agitation and alkylated with 20 mM iodoacetamide for 30 minutes at room temperature in the dark. Then, samples were acidified with phosphoric acid at a final concentration of 1.2 %. A ratio of 1:10 w: w Trypsin TPCK Treated (Worthington-Biochem) was used to digest the samples for 3 hours at 47 °C. Peptides were eluted sequentially with 50 mM TEAB, 0.2 % formic acid (FA) and 0.2 % FA in 50 % acetonitrile (MeCN). Eluted peptides were dried down and resuspended in loading buffer reverse phase (2 % MeCN, 0.1 % trifluoroacetic acid (TFA)).

### Data-independent acquisition mass spectrometry (DIA-MS)

Peptide samples from a 24-hour experiment were injected on a Dionex Ultimate 3000 RSLC (Thermo Fisher Scientific) connected to an Q Exactive Hybrid Quadrupole-Orbitrap mass spectrometer (Thermo Fisher Scientific). Samples were injected on a PepMap 100 C18 LC trap column (300 µm ID x 5 mm, 5 µm, 100 Å) followed by separation on an EASY-Spray column (50 cm x 75 µm ID, PepMap C18, 2 µm, 100 Å) (Thermo Fisher Scientific). Buffer A consisted of water containing 0.1 % FA and Buffer B of 80 % MeCN containing 0.1 % FA. Peptides were separated with a linear gradient of 3 − 35 % Buffer B over 120 minutes followed by a step from 35 − 90 % Buffer B in 0.5 min at 250 nL/min and held at 90 % for 4 min. The gradient was then decreased to 3 % Buffer B in 0.5 minutes at 250 nL/min for 20 minutes. Column temperature was controlled at 45°C. The Q Exactive Hybrid Quadrupole-Orbitrap mass spectrometer was operated in positive ion mode. MS scan spectra were acquired in the range of m/z 400 − 1,300 with a standard automatic gain control (AGC) target and a maximum injection time of 60 ms, at a resolution of 60,000. Targeted MS2 scan spectra were acquired in the range of m/z 400 − 1,000 using 16 m/z quadrupole isolation windows, AGC target of 1E6, at a resolution of 15,000, and higher-energy collision-induced dissociation (HCD) fragmentation was performed in one-step collision energy of 27 %. An electrospray voltage was static and capillary temperature of 275°C, with expected LC peak widths of 30 seconds. No sheath and auxiliary gas flow was used.

### Data-independent acquisition parallel accumulation, serial fragmentation (diaPASEF)

Solvent A was 0.1 % FA in water and Solvent B was 0.1 % FA in MeCN. All centrifugation steps were 700×g for 60 seconds. For peptide loading onto EvoTips (Evosep), tips were first activated by soaking in 1-propanol, then washed with 20 μL of Solvent B by centrifugation. Washed tips were conditioned by soaking in 1-propanol until the C18 material appeared pale white. Conditioned tips were equilibrated (Bruker) centrifugation with 20 μL Solvent A. Samples were then loaded into the tips while the tips were soaking in Solvent A to prevent drying, peptides were then bound to the C18 material by centrifugation. Tips were washed by centrifuging with 20 μL Solvent A. Next, 100 μL Solvent A was added to the tips and the tips were centrifuged at 700×g for 10 seconds. Samples were then immediately analysed by LC-MS/MS.

Peptide samples from 72-hour experiment were injected on an EvoSep One LC (Evosep) connected to a timsTOF HT mass spectrometer (Bruker) using a 15 cm Aurora Elite C18 column with integrated captive spray emitter (IonOpticks), at 50°C.

Buffer A was 0.1 % formic acid in HPLC water, buffer B was 0.1 % formic acid in acetonitrile. The pre-set Whisper100 20 samples per day method was used (the gradient was 0–35 % buffer B, 100 nl/min, for 58 minutes). The timsTOF HT was operated in DIA parallel accumulation, serial fragmentation (diaPASEF) mode. TIMS ion accumulation and ramp times were set to 100 ms, total cycle time was ∼1.8 seconds, and mass spectra were recorded from 300–1200 m/z. The mass spectrometer was operated in diaPASEF mode using variable width IM-m/z windows, designed using py_diAID (23). There were no overlap windows in m/z, but ion mobility windows were acquired with varying overlaps, as detailed in Table S1. Collision energy was applied in a linear fashion, with ion mobility of 0.6–1.6 Vs/cm^2^, and collision energy from 20 eV to 59 eV.

### MS data analysis

The DIA files were analysed library-free using DIA-NN (version 1.8) (24), searched against SwissProt Homo sapiens database with isoforms containing 42,393 sequences (downloaded on 14 March 2021), and contaminant FASTA from Ling Hao lab (25). The search parameters employed in DIA-NN were as follows: a minimum peptide length of 7 and a maximum of 30 amino acids, a precursor charge range of 1–4, a precursor m/z range of 300–1800, and a fragment ion m/z range of 200–1800. C-carbamidomethylation was designated as the fixed modification, while N-terminal M excision was defined as a variable modification. Trypsin specificity was set to two missed cleavages and a protein and PSM false discovery rate of 1 %, respectively. Intensity data were transformed (log 2) and filtered to contain at least two unique peptides and at least three valid values in one group for comparisons. The statistical analysis was performed using the R package Limma with a q-value (adjusted p-value, taking into account the false discovery rate (FDR)) threshold of FDR adjusted p-value < 0.05 (26). Perseus (version 2.0.11.0) (27) was used to generate the heatmaps. Kyoto Encyclopaedia of Genes and Genomes (KEGG) (28) was used to generate the pathway maps. Gene set enrichment analysis (GSEA) was performed using the GSEA-R, a Bioconductor implementation of GSEA from Broad Institute (29). Analysis was run with 10,000 permutations with Benjamini-Hochberg adjusted p-value threshold < 0.05. Normalised enrichment score and nominal P value were measured. Script used is available in the Supporting Information, R Script.

### Data availability statement

The mass spectrometry proteomics data have been deposited to the ProteomeXchange Consortium (30) via the PRIDE partner repository (31) with the data set identifier: PXD045276.

### Experimental Design and Statistical Rationale

MS-based proteomics experiments were conducted on K562, MOLT-4, NCI-H929, and U937 cell lines, each in three biological replicates to ensure robustness and reproducibility in capturing inherent biological variability within the experimental conditions. These cell lines were exposed to trilaciclib or vehicle (DMSO) for 24 hours. Additionally, K562 cells were exposed to trilaciclib, palbociclib, or vehicle (DMSO) for 72 hours. The replicates were processed collectively, and proteins were digested using trypsin, as previously described in ‘Sample preparation for LC-MS/MS.’ The resulting proteome samples underwent analysis using liquid chromatography-tandem mass spectrometry (LC-MS/MS). Proteins with significantly differential abundance between experimental conditions were identified by applying a student’s t-test with a cutoff of log2 fold change greater than ±0.58 and an adjusted p-value < 0.05. The sections ‘Sample preparation for LC-MS/MS,’ ‘Data-independent acquisition mass spectrometry (DIA-MS),’ ‘Data-independent acquisition parallel accumulation, serial fragmentation (diaPASEF),’ and ‘MS data analysis’ provide a comprehensive discussion of various sample preparation, MS parameters, and analysis parameters.

Validation of these results was performed in K562 and A549 cell lines, with four biological replicates for IC50 curves, beta-galactosidase activity, LysoTracker, proliferation, cell cycle, and Western blots experiments; three biological replicates for apoptosis assays; and six biological replicates for Seahorse experiments. The number of biological replicates is indicated in each figure caption. The statistical significance of the distinct experiments was determined after a normality test by using Student’s t-test, one-way ANOVA, or the Kruskal– Wallis’s test in GraphPad Prism (version 8).

## RESULTS

### The K562 cell line is resistant to trilaciclib-induced non-apoptotic cell death

Our goal was to delve into the impact of trilaciclib across various haematological cancers. We utilised the Cancer Dependency Map project data and specifically interrogated 694 cell lines which had undergone whole-genome CRISPR screening to identify dependencies. Since trilaciclib is a specific inhibitor of CDK4/6, which plays a crucial role in the transition from G1 to S phase, we explored the consequences of CRISPR knockout of *TP53*, *CDK4*, *CDK6*, and *RB1* genes on trilaciclib sensibility (**Figure 1A**). The absence of *CDK4* and *CDK6* resulted in reduced sensitivity to trilaciclib, as these are its main targets. However, CRISPR knockout of *TP53* and *RB1* genes heightened cell sensitivity to trilaciclib. We performed further analysis of the DepMap PRISM repurposing of trilaciclib sensitivity screen across tumour tissues (**Figure 1B**), notably in haematological cancers (**Figure 1C**). Interestingly, we observed that certain cell lines, such as K562, are less sensitive to trilaciclib, while others, such as U937 and JURKAT, exhibit greater sensitivity. To validate these findings, we examined different concentrations of trilaciclib using various cell lines derived from acute myeloid leukaemia (AML): U937; acute lymphoblastic leukaemia (ALL): MOLT-4 and JURKAT; multiple myeloma: H929; and from chronic myeloid leukaemia (CML): K562. After 72 hours of treatment, cell viability was measured and IC50 dose calculated for all of them: 0.72 µM for H929, 1.56 µM for MOLT-4, 1.77 µM for U937, and 2.97 µM for JURKAT (**Figure 1D**). However, we observed that K562 cells did not reach the 50 % mark of cell death. Indeed, more than 70 % of K562 cells were still alive with the highest dose tested here (10 µM), while the rest of the cell lines tested had less than 20 % of cells alive. Additionally, our findings revealed that trilaciclib increases late apoptotic or necrotic stages in all other tested cell lines (**Figure 1E**), which might potentially trigger an inflammatory response and lead to the recruitment of immune cells (32–34). Trilaciclib’s ability to enhance T-cell activation and upregulating major histocompatibility complex (MHC) class I and II, along with stabilising programmed death-ligand 1 (PD-L1) (35), and the activation of non-apoptotic cell death open up promising avenues for combination therapies with immune checkpoint inhibitors, which could enhance the effectiveness of cancer immunotherapy. This could create therapeutic vulnerabilities that can be exploited, as demonstrated with the combination of palbociclib and KIT inhibition in RUNX1/ETO AML with activated KIT proto-oncogene mutations (36). Furthermore, trilaciclib can protect healthy cells from chemotherapy-induced DNA damage, reducing the activity of caspases 3 and 7, and improving treatment outcomes (37). This protective effect on healthy cells is particularly relevant in haematological cancers, where aggressive chemotherapy regimens are often used.

**Figure 1.**
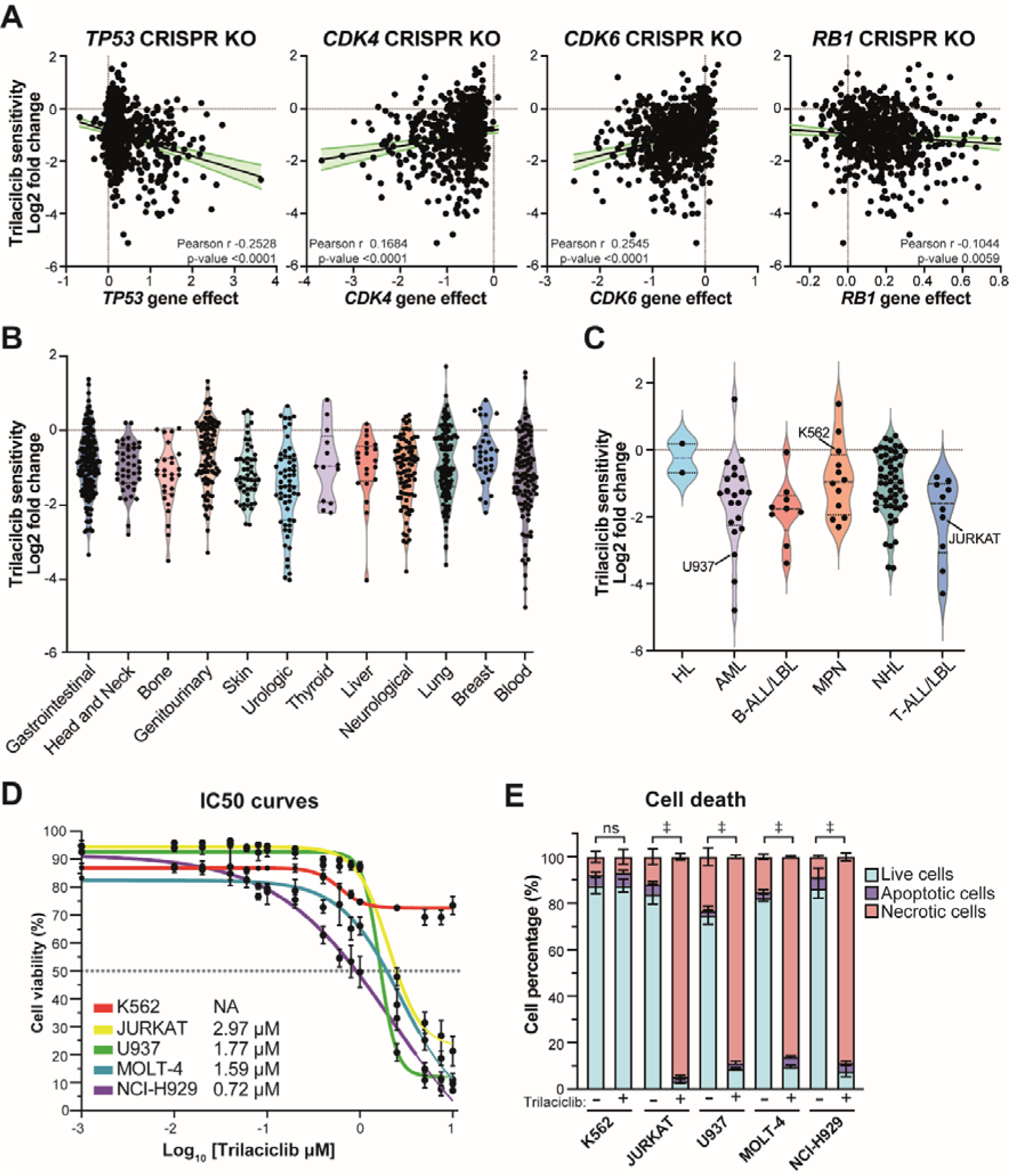
Trilaciclib induces cell death in a few cell lines, but not in K562, which exhibits resistance. **A)** The fitness effects of CRISPR-Cas9 gene knockout on *TP53*, *CDK4*, *CDK6*, and *RB1* genes (DepMap Public+Score, Chronos, https://depmap.org/portal/)) were correlated with trilaciclib sensitivity across 694 cell lines. Pearson correlation coefficient (r) and p-value are shown in each graph. **B-C)** The sensitivity of trilaciclib varies across different histologic cancer types (DepMap, BRD: BRD-K00003412-300-01-9, PRISM Repurposing Public 23Q2; https://depmap.org/portal/). In **C)** blood cancers are subdivided into Hodgkin lymphoma (HL, 2 cell lines), acute myeloid leukaemia (AML, 21 cell lines), T-cell acute lymphoblastic leukaemia/ lymphoma (T-ALL/LBL, 10 cell lines), myeloproliferative neoplasms (MPN, 12 cell lines), non-Hodgkin lymphoma (NHL, 54 cell lines), and B-cell acute lymphoblastic leukaemia/ lymphoma (B-ALL/LBL, 9 cell lines). **D)** The inhibition concentration at 50 % cell death (IC50) curves were analysed in K562, JURKAT, U937, MOLT-4, and NCI-H929 cells after 72 hours of treatment. Standard deviation of four biological replicates is shown. **E)** Percentage of live cells (Annexin V negative and 7-AAD negative cells, light blue), apoptotic cells (Annexin V positive and 7-AAD negative cells, purple), and necrotic cells (7-AAD positive cells, pink) after 72 hours of treatment. The statistical significance of the comparisons with resting is indicated as follows: ‡, P ≤ 0.001; ns, not significant. Standard deviation of three biological replicates is shown.

### Trilaciclib impairs cell cycle progression and induces autophagy

Trilaciclib is a CDK4/6 inhibitor used in lung cancer as a myeloprotective agent. Its mechanism involves transiently impeding the cell cycle progression of haematopoietic stem cells, enabling them to enter a protective quiescent state. To decipher the mechanism by which trilaciclib can induce non-apoptotic cell death in certain cell lines but not in K562, we employed mass spectrometry-based proteomics to characterise the effect of short-term, 24-hour trilaciclib treatment. Using the QE mass spectrometer in data-independent acquisition (DIA) mode, we detected 5655 proteins using a library free analysis in trilaciclib-treated K562 cells for 24 hours (**Figure S1A** and **Table S2**). Proteomics analysis revealed the ability of trilaciclib to stabilise CDK4 and CDK6 while downregulating proteins associated with cell cycle progression and proliferation such as CDK1/2, RB1, AURKA/B, BUB1, PLK1, CDC45, CCNB1, CDC20, or MKI67 (**Figure 2A**). At 24 hours of trilaciclib treatment, we noted an upregulation of CDK4/6 proteins. Subsequently, we conducted a Western blot-based Cellular Thermal Shift Assay (CETSA) to validate the stability of CDK4 and CDK6 during their interaction with trilaciclib. The stabilisation was more pronounced for CDK4 than for CDK6 (**Figure S2**), corresponding to a higher upregulation of CDK4. Additionally, proteomic analysis of the effects of trilaciclib was conducted on U937, MOLT-4, and NCI-H929 cells, resulting in the identification and quantification of 5639, 5850, and 5833 proteins, respectively (**Figure S1B-C** and **Table S3-S5**). Despite their differing sensitivities to trilaciclib, these cell lines – U937, MOLT-4, NCI-H929, and K562 – exhibited alterations in cell cycle-related proteins. These findings underscore the potential of trilaciclib as a broad-spectrum CDK4/6 inhibitor that can impact haematological cancer cells’ growth and proliferation. Notably, they shared 46 interconnected proteins (**Figure 2B-C**), of which 32 appeared to be involved in cell cycle regulation (**Figure 2C**). GSEA analysis of K562 cells revealed downregulated protein sets enriched in processes associated with the cell cycle, G2/M checkpoint, and E2F targets (**Figure S3**). Nevertheless, K562 is resistant to trilaciclib; therefore, we also investigated the upregulated proteins. Interestingly, they do not share any common proteins, with only ITPR1 (inositol 1,4,5-trisphosphate receptor type 1) being significantly upregulated in all cell lines except K562 (**Figure 2D**). ITPR1 is involved in diverse cellular functions such as calcium signalling and is associated with cell death, cell cycle, autophagy, senescence, and lipid metabolism (37–39). Furthermore, in K562, U937, and MOLT-4 cells, processes associated with lysosomes and lipid biosynthesis/metabolism exhibited upregulation, which is intricately controlled during senescence and cell death (40, 41). Therefore, trilaciclib might have direct and common targets that impact the cell cycle such as CDK4/6, but it exhibits cell-line specificity when it comes to lipid metabolism and associated cellular processes, including senescence or cell death.

**Figure 2.**
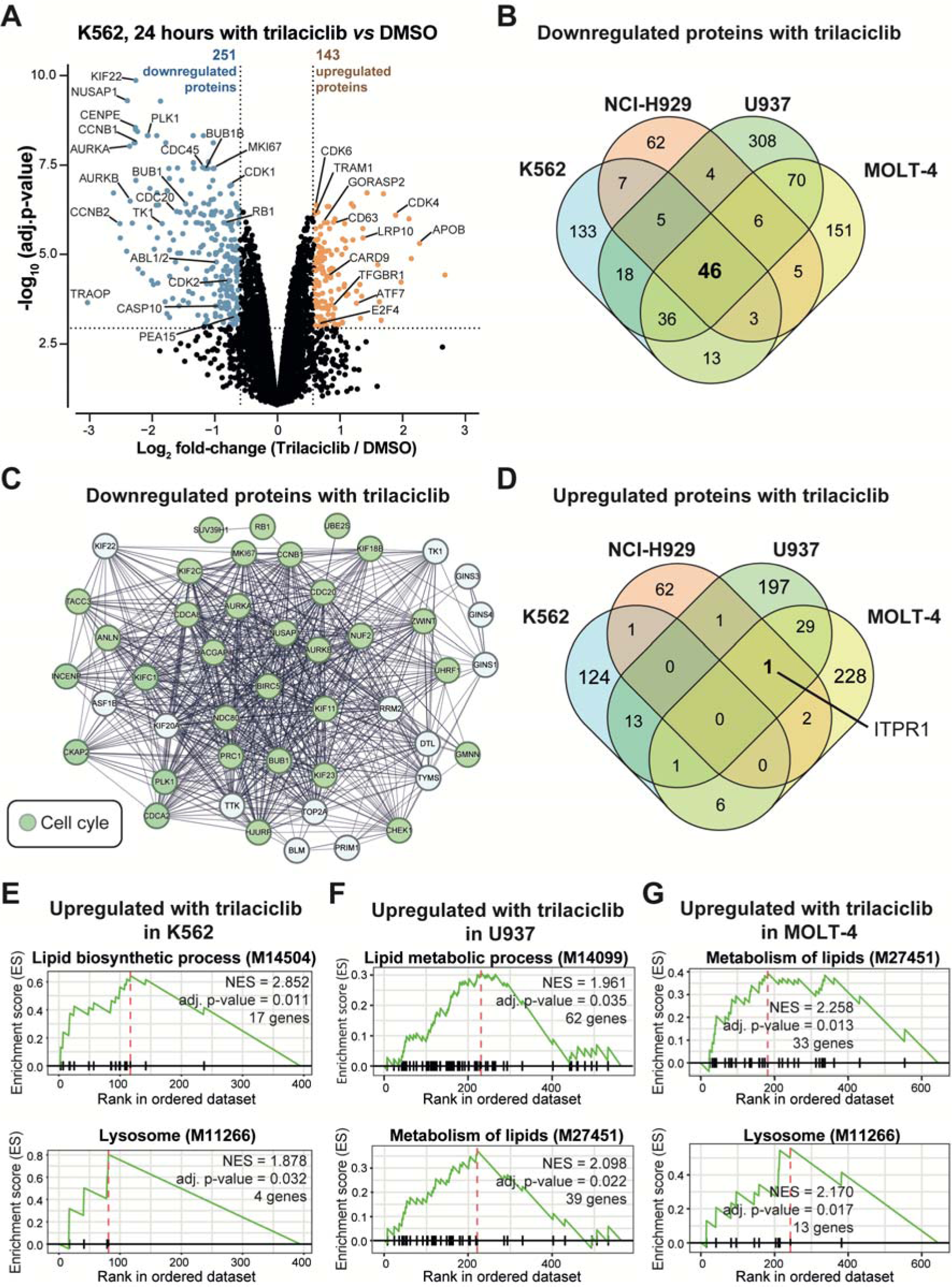
Trilaciclib induces cell cycle arrest and alters lipid biosynthesis and metabolism. **A)** The volcano plot illustrates the differentially expressed proteins between trilaciclib- and DMSO-treated K562 cells after 24 hours. Significant upregulated proteins are shown in orange, while downregulated proteins are shown in blue (FDR adjusted p-value < 0.05). **B)** Venn diagram illustrates the significant downregulated proteins in trilaciclib-treated K562, U937, MOLT-4, and NCI-H929 cell lines. **C)** STRING analyses reveal the 46 common significant downregulated proteins shared in trilaciclib-treated K562, U937, MOLT-4, and NCI-H929 cell lines. The proteins related to cell cycle regulation are highlighted in green. **D)** Venn diagram illustrates the significant upregulated proteins in trilaciclib-treated K562, U937, MOLT-4, and NCI-H929 cell lines. **E-G)** GSEA analysis shows significant enrichment in upregulated signalling pathways after 24 hours of trilaciclib treatment compared to DMSO in **E)** K562, **F)** U937, and **G)** MOLT-4 cells.

Unlike other cellular processes, senescence is not an immediate response and may take days (42). To investigate this further, we conducted a proteomics study after 72 hours of treatment with trilaciclib in K562 cells, which was analysed using the TimsTOF HT mass spectrometer in diaPASEF mode (**Figure S4** and **Table S6**). We detected 7404 proteins using a library free analysis in DIA-NN. Proteomics analysis conducted after a 72-hour treatment with trilaciclib also identified stabilised CDK4 and CDK6 and downregulated proteins associated with cell cycle progression and proliferation such as CDK1/2, RB1, AURKB, BUB1, PCNA, TK1, or MKI67 (**Figure 3A**). Following a 72-hour treatment with trilaciclib, proteins such as CD63, cathepsin A and D (CTSA and CTSD), renin (REN), beta-galactosidase (GLB1), and LAMP1 were upregulated. A comprehensive analysis of the dysregulated proteins by trilaciclid treatment across all cell types examined here unveiled an enrichment of upregulated proteins in K562 cells, particularly evident at 72 hours post-treatment, which correlates with enhanced lysosomal activity and autophagy (**Figure 3B**). Conversely, proteins associated with apoptosis and ferroptosis exhibited decreased expression in all trilaciclib-treated cells, whereas those involved in the negative regulation, such as cIAP and XIAP, were upregulated (**Figure 3B**). Furthermore, trilaciclib stimulated the metabolism of lipids, lysosomes, and secretion, which might trigger senescence (**Figure 3C**). In contrast, processes associated with the cell cycle, RB1, and E2F targets were downregulated at 72 hours of trilaciclib treatment in K562 cells compared to DMSO (**Figure 3C**). Additionally, we incorporated palbociclib treatment into the study, given its ability to induce senescence and its role as a CDK4/6 inhibitor (43–45). Palbociclib, another CDK4/6 inhibitor, also induced senescence and reduced proliferation in K562 cells, but it did not exhibit the same dysregulated proteins, indicating that trilaciclib operates through a different molecular mechanism to induce senescence. There were upregulated proteins only with trilaciclib treatment involved in secretion and extracellular vesicles, such as renin (REN), CD63, tetraspanin-3 (TSPAN3), GM2A, ITM2B, DHCR24, or MME (**Figure S5A**). Specifically, the tetraspanin CD63, upregulated in response to trilaciclib treatment compared with palbociclib, was associated with the regulation of cellular senescence, controlling endocytic trafficking and exosome formation (46). In addition, proteins related to the endomembrane system, endoplasmic reticulum, and Golgi vesicle-mediated transport, such as RAB43, SEC24D, SEC11C, SLC33A1, or SLC6A19, were upregulated with trilaciclib compared with palbociclib (**Figure S5B**).

**Figure 3.**
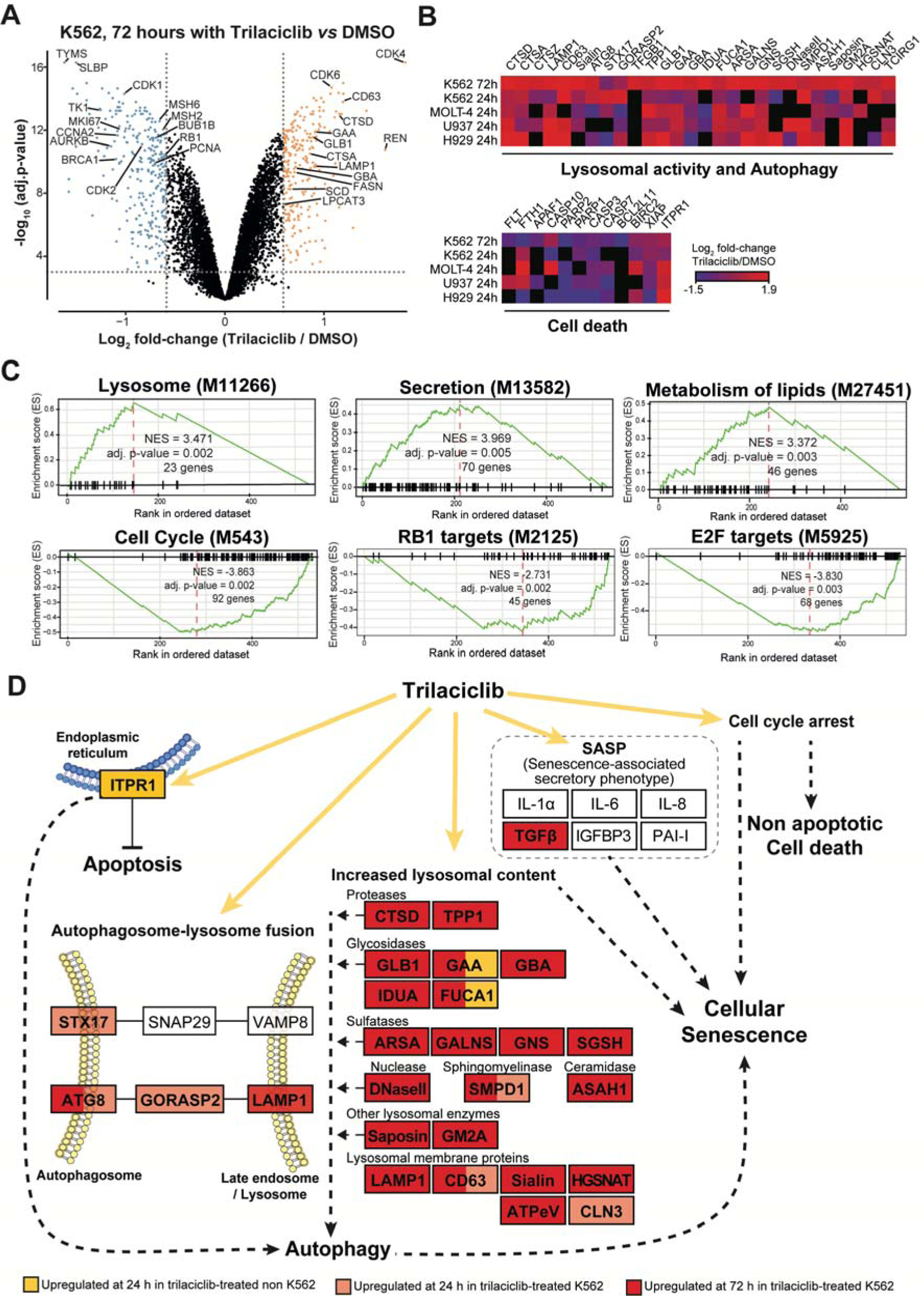
Trilaciclib arrests cell cycle, increases lysosome activity and autophagy in K562 cells. **A)** Volcano plot illustrates the differentially expressed proteins between trilaciclib- and DMSO-treated K562 cells after 72 hours. Upregulated proteins are shown in orange, while downregulated proteins are shown in blue (FDR adjusted p-value < 0.05). **B)** The heatmap represents the log2 fold-changes between trilaciclib- and DMSO-treated cells independent of adjusted p-value. **C)** GSEA analyses show significant enrichment in trilaciclib-treated cells compared to DMSO. **D)** KEGG pathway diagram illustrating the mechanism by which trilaciclib induces cell death and senescence in K562 compared to U937, MOLT-4, and NCI-H929 cell lines. Significant upregulated proteins (FDR adjusted p-value < 0.05) are shown at 24 hours in at least two of the trilaciclib-*versus* DMSO-treated U937, MOLT-4, and NCI-H929 cell lines (orange), at 24 hours in trilaciclib-*versus* DMSO-treated K562 (light red), and at 72 hours in trilaciclib-*versus* DMSO-treated K562 (dark red).

Overall, this proteomics analysis of trilaciclib treatment across various cell types reveals intriguing insights into its dual role. Firstly, in all the cells lines treated with trilaciclib, there is evidence of cell cycle disruption, manifested by cell cycle arrest and non-apoptotic cell death except in K562. Moreover, cells where trilaciclib induces non-apoptotic cell death, only ITPR1 exhibited upregulation, accompanied by low levels of autophagic proteins. This suggests that ITPR1 might play a crucial role in the mechanism by which trilaciclib impacts the apoptosis and triggers autophagy. Specifically, the enrichment of upregulated proteins in K562 cells post-trilaciclib treatment, suggests a correlation with enhanced lysosomal activity, autophagy, and cell cycle arrest (**Figure 3D**). This highlights a potential mechanism through which trilaciclib induces senescence in K562 cells. Notably, trilaciclib operates through distinct mechanisms compared to palbociclib, as evidenced by the differing dysregulated protein profiles. This suggests a unique molecular signature by which trilaciclib induces senescence.

### Trilaciclib induces senescence in K562 (CML) and A549 cells (NSCLC)

To validate the observed changes in the proteome, we first investigated beta-galactosidase and lysosomal activity. The findings revealed a significant increase in beta-galactosidase activity in trilaciclib-treated K562 and A549 cells (**Figure 4A** and **Figure S6A**), indicating senescence induction. Moreover, the alteration in lysosomal activity highlights the involvement of lysosomes in autophagy and senescence (**Figure 4B** and **Figure S6B**). Next, we examined the proliferation rate of trilaciclib-treated K562 cells for 12 days in K562 cells (**Figure 4C**) and for 7 days in A549 cells (**Figure S6C**). Consistent with previous findings (47), palbociclib showed a reduction in proliferation (**Figure 4C** and **Figure S6C**). Trilaciclib effectively blocked proliferation in K562 cells without affecting the cell viability (**Figure 1E**). Further analysis of the cell cycle phases revealed that both trilaciclib and palbociclib arrested the cells in the G0/G1 phase (**Figure 4D** and **Figure S6D**). Trilaciclib-induced cell cycle arrest was independent of p21 and p53 and mediated through the inhibition of CDK4/6, as K562 cells are deficient in p53 (48), affecting the phosphorylation of the retinoblastoma protein (p-RB) (**Figure 4E** and **Figure S6E**). Trilaciclib demonstrated potential as a treatment by selectively inhibiting CDK4, CDK6, and CDK5, leading to a cell cycle arrest (37). Thus, trilaciclib disrupts the normal cell cycle machinery in K562 and A549 cells, resulting in the inhibition of cell proliferation. This property may be particularly beneficial in haematological cancers where uncontrolled cell proliferation is a hallmark. It is important to note that the proteomics data presented is specific to K562 cells. Nevertheless, these proteomic alterations not only contribute to trilaciclib’s observed capacity to induce senescence in K562 cells but also hold significance in the induction of senescence in A549 cells. These discoveries provided insights into the mechanisms by which trilaciclib exerts its effects and emphasise the potential therapeutic relevance of trilaciclib across diverse cell types in cancer treatment.

**Figure 4.**
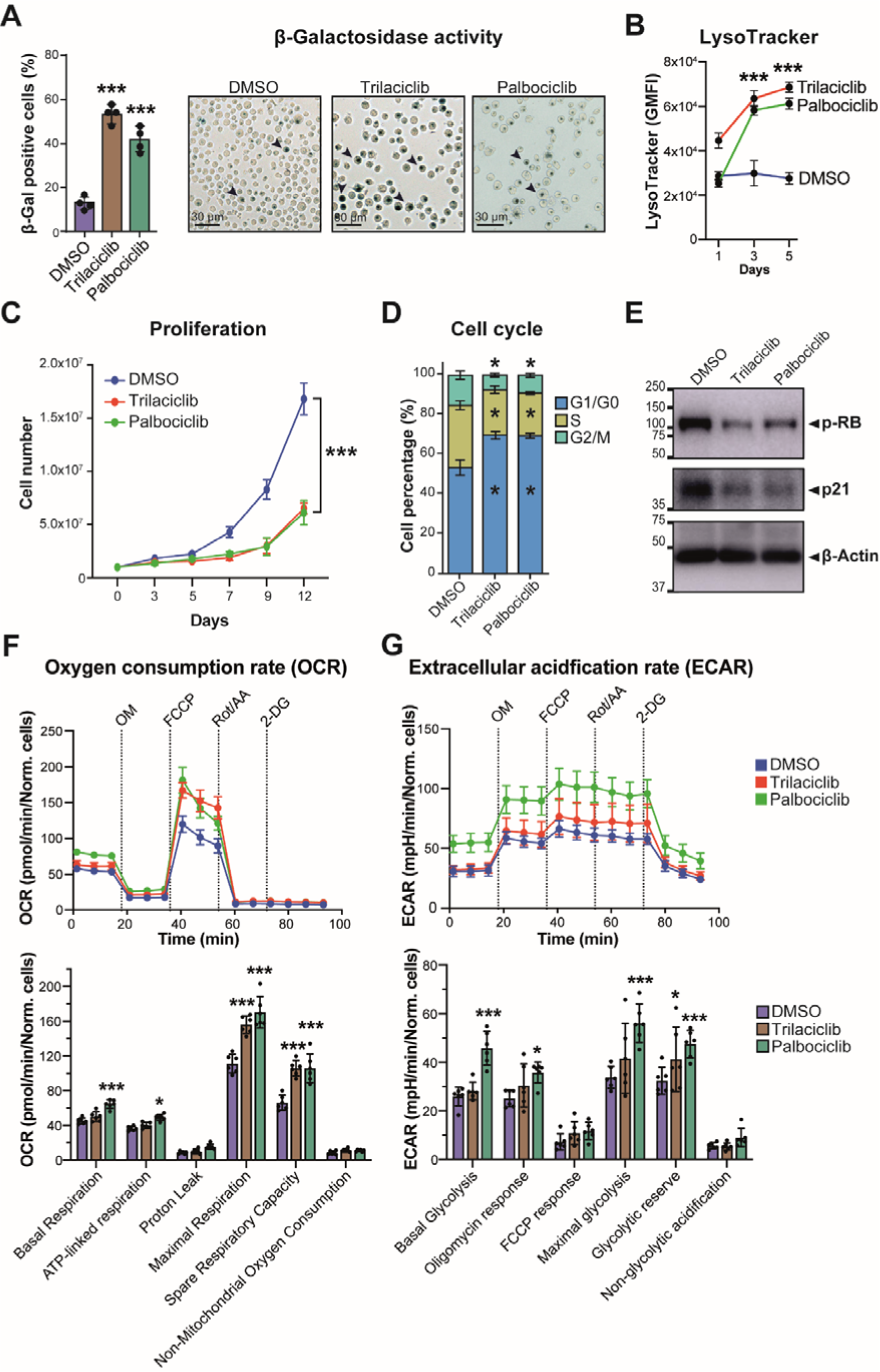
Trilaciclib induces senescence in K562 cells. **A)** Beta-galactosidase activity and **B)** LysoTracker were measured in trilaciclib-, palbociclib- and DMSO-treated K562 cells after 72 hours. **C)** K562 cells were counted using trypan blue staining for 12 days to assess viable cells and compared to the number of cells seeded at initial time (t_0_). **D-E)** K562 cells were treated with trilaciclib, palbociclib or vehicle (DMSO) for 5 days. Subsequently: **D)** cell cycle analysis was performed by staining cells with PI/RNAse buffer and analysing them by flow cytometry; and **E)** Western blot analysis was conducted to examine protein levels of senesce markers. The standard deviation of four biological replicates is shown. A representative image of four replicates is shown. Relative mobilities of reference proteins (masses in kDa) are shown on the left of each blot. **F)** Oxygen consumption rate (OCR) and **G)** extracellular acidification rate (ECAR) in trilaciclib-, palbociclib- and DMSO-treated K562 cells after 72 hours. The statistical significance of the comparisons with resting is indicated as follows: ***, P ≤ 0.001; *, P ≤ 0.05. Standard deviation represents biological replicates.

Senescent cells often undergo metabolic reprogramming. Dysfunctional mitochondria can generate increased levels of reactive oxygen species (ROS), which can contribute to oxidative stress and cellular damage, accelerating the senescence process (49). Effect that we can see after 3 days of treatment with trilaciclib (**Figure 4F-G**). However, senescent cells also rely less on oxidative phosphorylation for ATP production, leading to a reduction in oxygen consumption rate (OCR), favouring glycolysis over oxidative phosphorylation. Consequently, senescent cells exhibit an increased ECAR after 5 days with trilaciclib, indicating enhanced glycolytic activity (**Figure S7**). However, senescent cells also exhibited a shift towards glycolysis over oxidative phosphorylation. This metabolic shift, known as the Warburg effect (50), allows senescent cells to sustain ATP levels and adapt to their altered metabolic state (51). When tumour cells encounter stress, such as from therapy, autophagy releases metabolic precursors for ATP generation and facilitates immune evasion (18). Trilaciclib-treated cells, especially after several days of exposure, showed signs of metabolic changes, i.e., heightened glycolytic activity.

Overall, our study reveals intriguing insights into the effects of trilaciclib on various cell lines. Trilaciclib demonstrates a remarkable capability to induce cell death in cell lines derived from AML, ALL, and myeloma, while concurrently eliciting senescence in the K562 cells. The publicly available DepMap database was used to determine sensitivity factors across human cells. Notably, when we compared trilaciclib with palbociclib, another CDK4/6 inhibitor, we observed an interesting pattern. While palbociclib exhibited a high correlation with the effects induced by trilaciclib, it showed lower overall sensitivity (**Figure 5A**). This observation suggests that trilaciclib may possess unique attributes that make it more effective. Intriguingly, we observed similar trends with TKIs, we explored the impact of imatinib and asciminib, representing the first-generation and latest-generation TKIs, respectively (**Figure 6B-C**). This finding suggests that the effects of trilaciclib might synergize or complement those of various TKIs. Furthermore, asciminib is effective in chronic-phase CML patients who have undergone extensive prior TKI treatment and have experienced treatment failure (8). This insight encourages further exploration of tailored combination therapies involving trilaciclib and TKIs, offering new prospects for improved treatment strategies in various cancer contexts.

**Figure 5.**
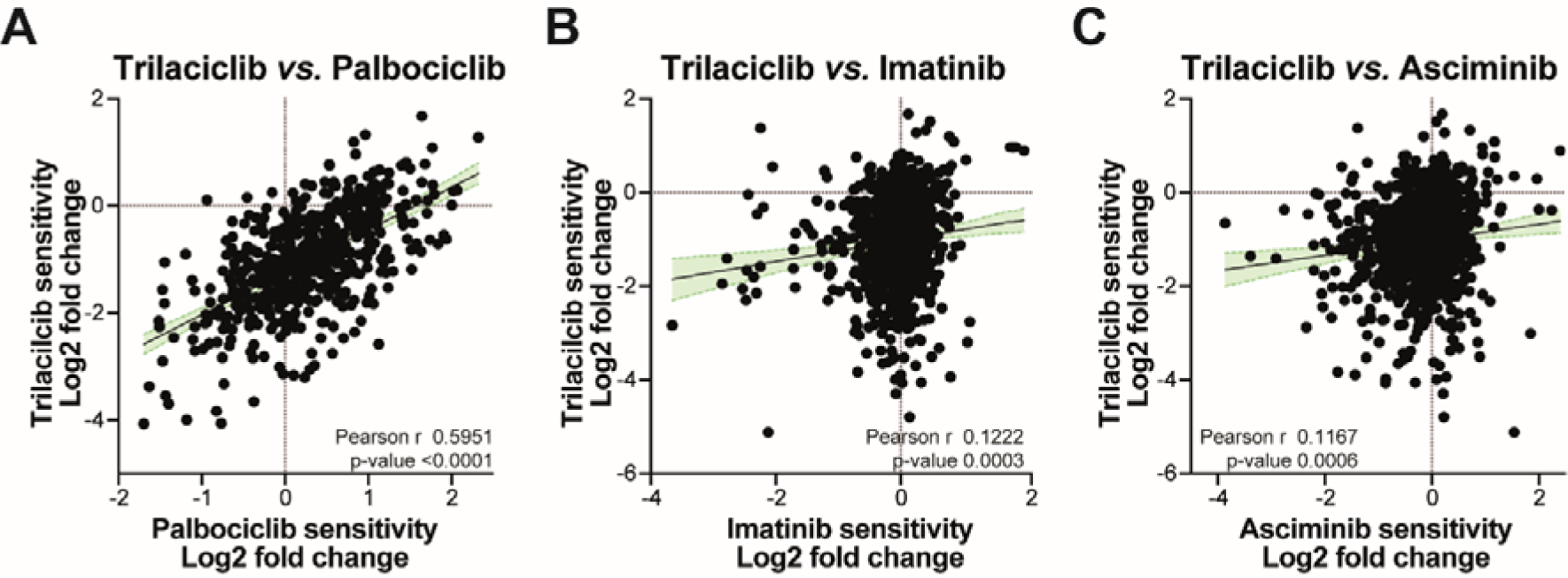
Trilaciclib exhibits higher sensitivity compared to palbociclib, imatinib, and asciminib. Sensitivity of trilaciclib were correlated with **A)** palbociclib, **B)** imatinib, and **C)** asciminib sensitivity across 539, 874, and 866 cell lines, respectively (DepMap, BRD: BRD-K00003412-300-01-9, BRD: BRD-K92723993-001-17-4, BRD: BRD-K00003456-001-01-9, and BRD: BRD-K51313569-003-03-3, PRISM Repurposing Public 23Q2; accessible at https://depmap.org/portal/). Pearson correlation coefficient (r) and p-value are shown in each graph.

## CONCLUSIONS

The administration of trilaciclib in cancer patients can trigger various outcomes. While it can lead to myelosuppression in healthy cells, it also instigates senescence in tumoral cells. These effects exemplify different facets of its pharmacological activity, particularly relevant in the treatment of NSCLC. Trilaciclib might be a therapeutic option for haematological cancers, particularly in cases where senescence induction may be beneficial, such as CML. Trilaciclib’s ability to arrest the cell cycle, inhibit proliferation, promote autophagy, and induce senescence suggests its versatility as a treatment strategy for various cancer types. Considering the potential drawbacks of senescence induction, such as chronic inflammation and tissue damage, it becomes essential to carefully balance the use of trilaciclib in managing haematological cancers. Additionally, the combination of CDK4/6 inhibitors like trilaciclib with senolytic compounds may offer new therapeutic strategies to address the negative effects of senescence and improve treatment outcomes. Further research and clinical trials are needed to validate these findings and explore trilaciclib’s full potential in haematological cancers and other cancer types, including non-small cell lung cancer.

## Abbreviations

AML: acute myeloid leukaemia
ALL: acute lymphoblastic leukaemia
CDK: cyclin-dependent kinase
CETSA: cellular thermal shift assay
CML: chronic myeloid leukaemia
DIA: data-independent acquisition
ECAR: extracellular acidification rate
ES-SCLC: extensive-stage small cell lung cancer
GSEA: gene set enrichment analysis
MS: mass spectrometry
HL: Hodgkin lymphoma
NSCLC: non-small cell lung carcinoma
T-ALL: T-cell lymphoblastic leukaemia
TKIs: tyrosine kinase inhibitors
OCR: oxygen consumption rate.

## Authorship contributions

M.H.-T.: formal analysis, methodology, validation, writing-original draft, review and editing, and visualisation. M.X.H.-Y.: formal analysis, methodology, validation, writing-original draft, and visualisation. A.M.F.: methodology. A.L.G.: methodology. A.D.: methodology. M.T.: resources, review and editing, and funding acquisition. J.L.M.-R.: conceptualization, methodology, validation, formal analysis, visualisation, review and editing, supervision, project administration, and funding acquisition. All authors provided critical feedback and helped shape the research, analysis, and manuscript.

## Notes

The authors declare no competing financial interest.

## ACKNOWLEDGEMENTS

This work was partly funded by a Welcome Trust Investigator Award (215542/Z/19/Z) to M.T. This research was partly funded by Newcastle Wellcome Trust Translational Partnership to J.L.M.-R. and M.T. M.H-T and M.X.H-Y were hosted by J.L.M-R through the Winter Studentship awarded by the Spanish Researchers in the United Kingdom Society (SRUK/CERU).

