## Supporting Information for "Proteomic Analysis Reveals Trilaciclib-Induced Senescence"

### CONTENT

**Figure S1. Proteomics quality controls for trilaciclib- and DMSO-treated cells for 24 hours.**

**Figure S2. Trilaciclib stabilises CDK4 and CDK6 in the K562 cell.**

**Figure S3. Trilaciclib induces cell cycle arrest in K562 cells.**

**Figure S4. Proteomics quality controls for trilaciclib-, palbociclib- and DMSO-treated K562 cells for 72 hours.**

**Figure S5. Trilaciclib and palbociclib trigger senescence through distinct mechanisms.**

**Figure S6. Trilaciclib induces senescence in A549 cells.**

**Figure S7. Trilaciclib enhances the Warburg effect in A549 cells.**

**R Script.**

**Table S1. Parameters for diaPASEF acquisition on the timsTOF HT mass spectrometer.** Table indicates the number of windows, mass/charge (m/z), and ion mobility (1/K0).

**Table S2. Proteomics ananlysis of trilaciclib-treated K562 cells for 24 hours.** Document indicates description, data, downregulated and upregulated gene ontology and KEGG enrichment analyses.

**Table S3. Proteomics ananlysis of trilaciclib-treated U937 cells for 24 hours.** Document indicates description, data, downregulated and upregulated gene ontology and KEGG enrichment analyses.

**Table S4. Proteomics ananlysis of trilaciclib-treated MOLT-4 cells for 24 hours.** Document indicates description, data, downregulated and upregulated gene ontology and KEGG enrichment analyses.

**Table S5. Proteomics ananlysis of trilaciclib-treated NCI-H929 cells for 24 hours.** Document indicates description, data, downregulated and upregulated gene ontology and KEGG enrichment analyses.

**Table S6. Proteomics ananlysis of trilaciclib-treated K562 cells for 72 hours.** Document indicates description, data, downregulated and upregulated gene ontology and KEGG enrichment analyses.

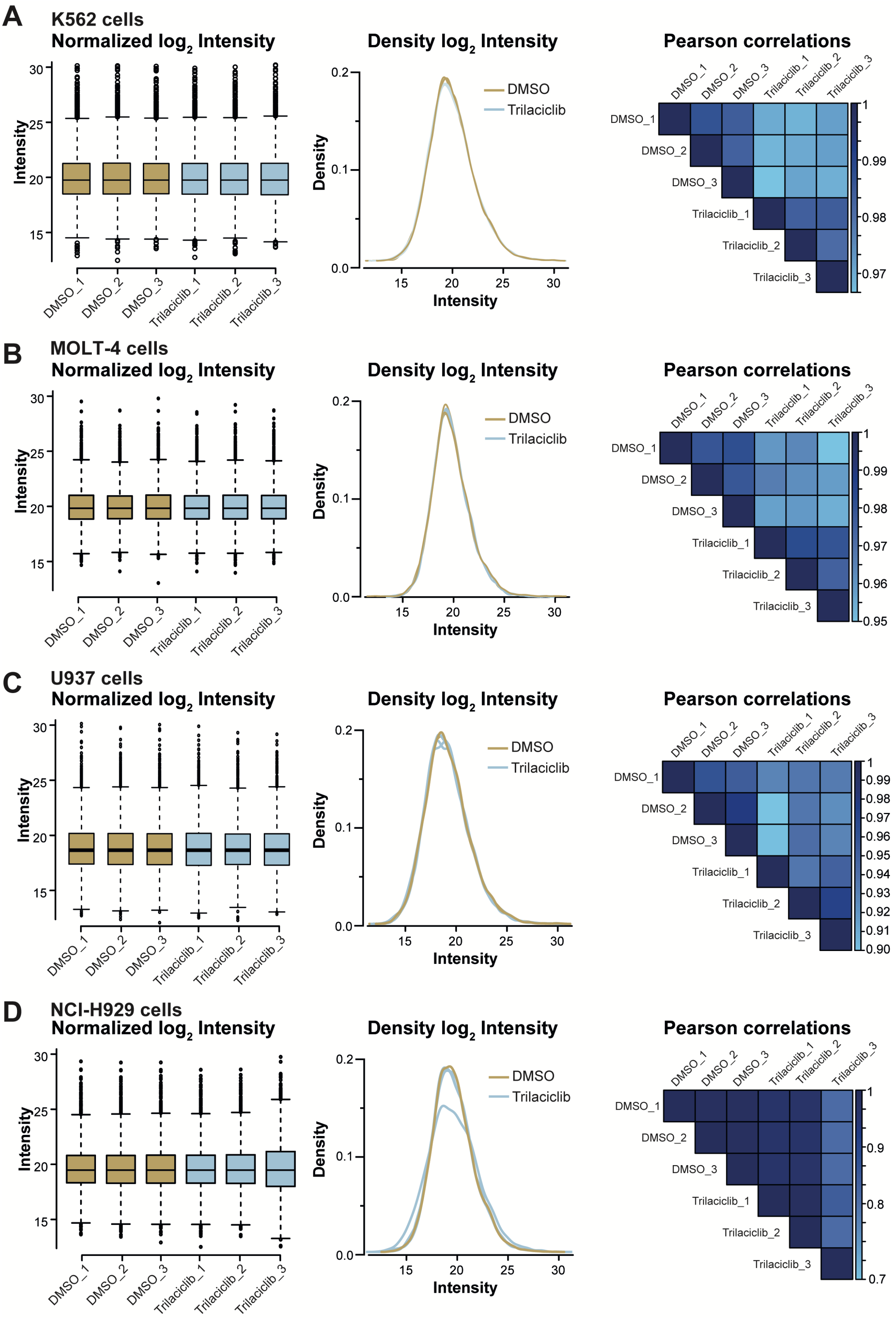
A
K562 cells
Normalized log2 Intensity
Intensity
DMSO_1
DMSO_2
DMSO_3
Trilaciclib_1
Trilaciclib_2
Trilaciclib_3
Density log2 Intensity
Density
DMSO
Trilaciclib
Intensity
Pearson correlations
B
MOLT-4 cells
Normalized log2 Intensity
Density log2 Intensity
Pearson correlations
C
U937 cells
Normalized log2 Intensity
Density log2 Intensity
Pearson correlations
D
NCI-H929 cells
Normalized log2 Intensity
Density log2 Intensity
Pearson correlations

**Figure S1. Proteomics quality controls for trilaciclib- and DMSO-treated cells for 24 hours. A)** K562, **B)** MOLT-4, **C)** U937, and **D)** NCI-H929 cells**.** Box and whisker plot and Kernel density plot show the $\log_2$ transformed normalised intensities of three biological replicates per group. The correlation matrix displays the Pearson correlation coefficients between $\log_2$ transformed intensities. The colour code reflects the correlation coefficient values.

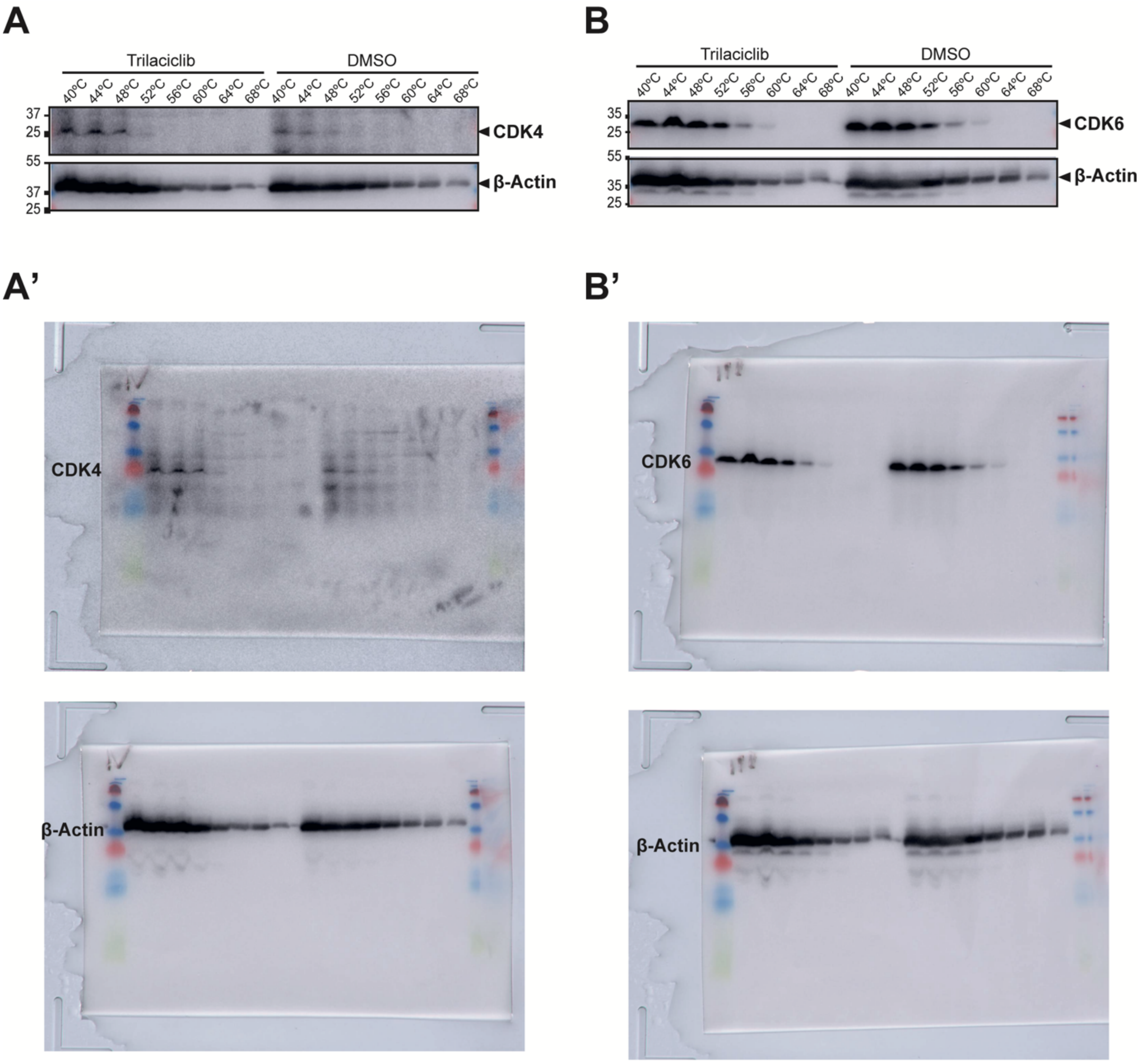


**Figure S2. Trilaciclib stabilises CDK4 and CDK6 in the K562 cell.** Determination of the thermostability of **A)** CDK4 and **B)** CDK6 at the indicated temperatures in trilaciclib- and DMSO-treated K562 cells for 1 hour. Beta-actin served as a loading control. A representative image of three replicates is shown. Relative mobilities of reference proteins (masses in kDa) are shown on the left of each blot. Full western blot membranes are shown in **A')** and **B')**.

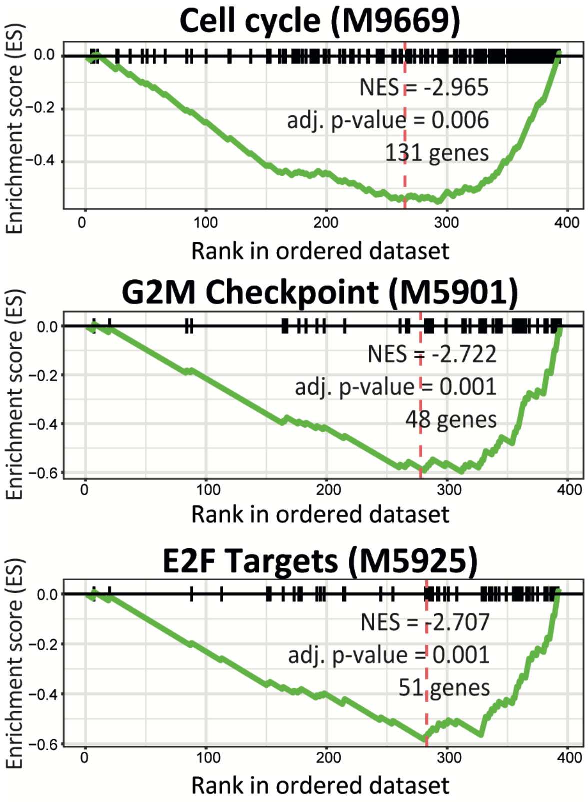


**Figure S3. Trilaciclib induces cell cycle arrest in K562 cells**. GSEA analyses show significant enrichment in downregulated proteins after 24 hours with trilaciclib compared to DMSO-treated cells.

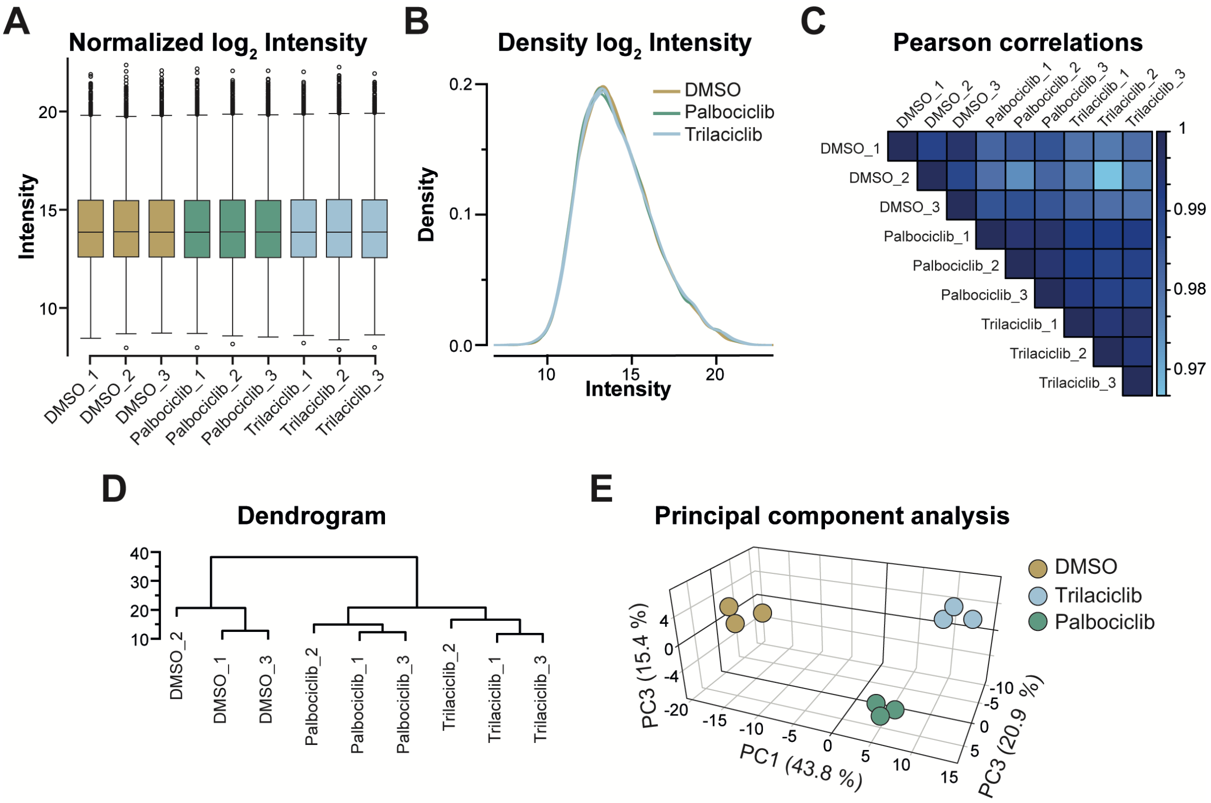


**Figure S4. Proteomics quality controls for trilaciclib-, palbociclib- and DMSO-treated K562 cells for 72 hours. A)** Box and whisker plot and **B)** Kernel density plot show the $log_2$ transformed normalised intensities of three biological replicates per group. **C)** the correlation matrix displays the Pearson correlation coefficients between $log_2$ transformed intensities. The colour code reflects the correlation coefficient values. **D)** Dendrogram illustrates the hierarchical clustering of samples. **E)** Principal component analysis (PCA) of the whole proteome dataset shows the three main principal components (PC).

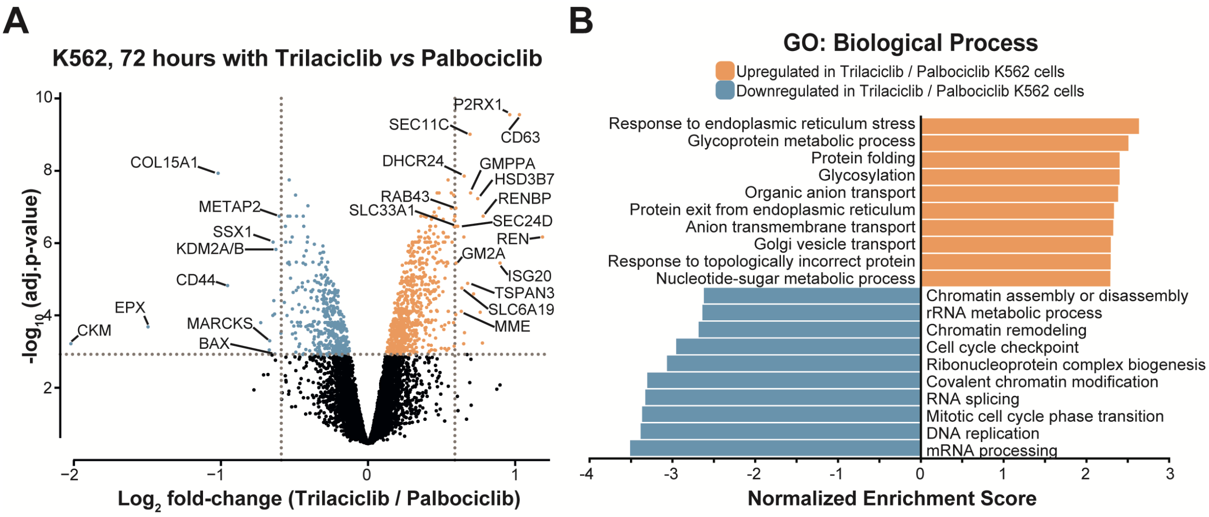


**Figure S5. Trilaciclib and palbociclib trigger senescence through distinct mechanisms.** **A)** Volcano plot illustrates the differentially expressed proteins between trilaciclib- and palbociclib-treated K562 cells after 72 hours. Upregulated proteins are shown in orange, while downregulated proteins are shown in blue (FDR adjusted p-value < 0.05). **B)** Gene ontology (GO) analyses show the significant enrichment (FDR < 0.05) of biological processes in trilaciclib-treated cells compared to palbociclib-treated cells.

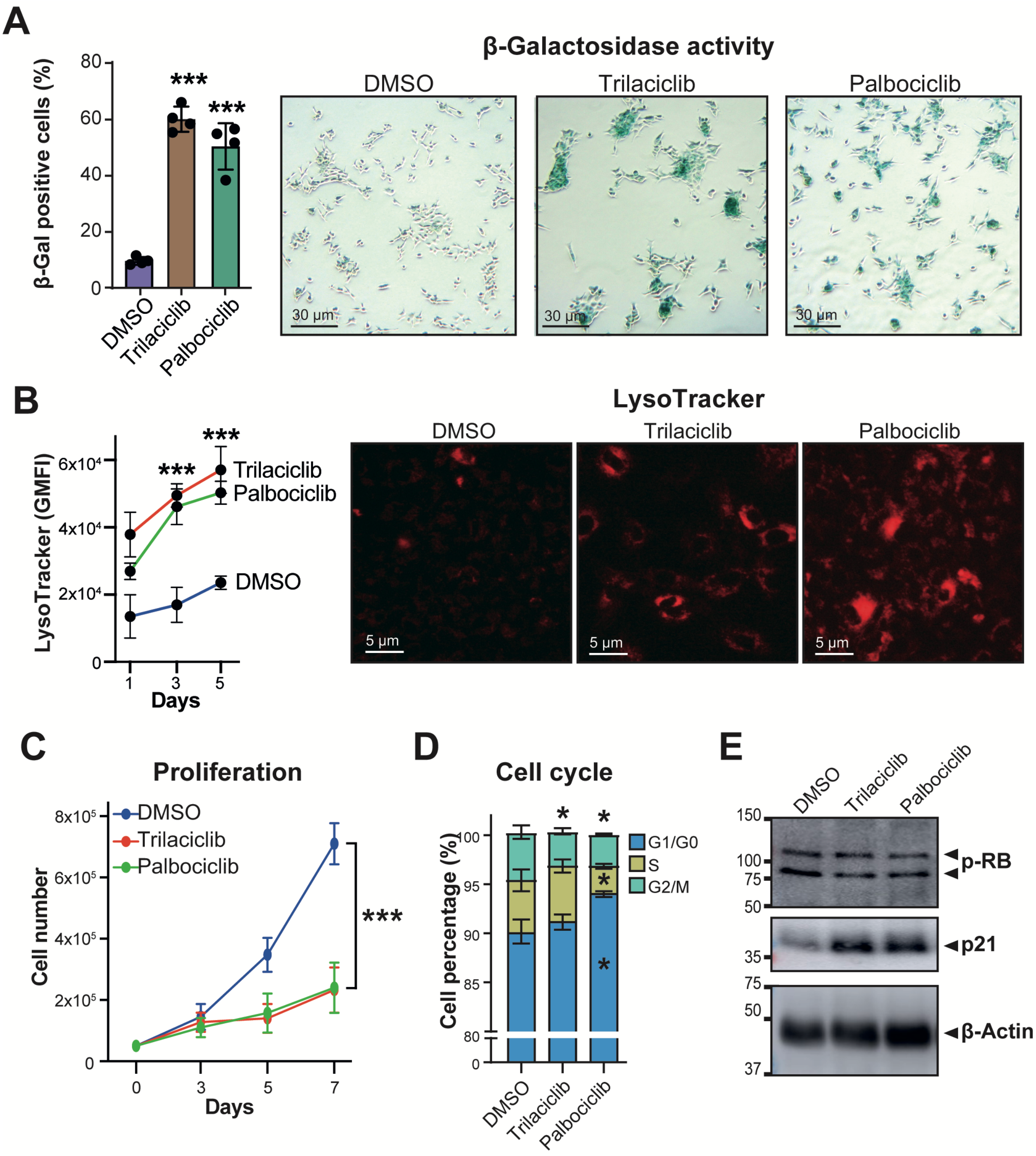


**Figure S6. Trilaciclib induces senescence in A549 cells. A)** Beta-galactosidase activity and **B)** LysoTracker were measured in trilaciclib-, palbociclib- and DMSO-treated A549 cells after 72 hours. **C)** A549 cells were counted using trypan blue staining for 7 days to assess viable cells and compared to the number of cells seeded at initial time ($t_0$). **D-E)** A549 cells were treated with trilaciclib, palbociclib or vehicle (DMSO) for 5 days. Subsequently: **D)** cell cycle analysis was performed by staining cells with PI/RNAse buffer and analysing them by flow cytometry; and **E)** Western blot analysis was conducted to examine protein levels of senesce markers. The standard deviation of four biological replicates is shown. A representative image of four replicates is shown. Relative mobilities of reference proteins (masses in kDa) are shown on the left of each blot. The statistical significance of the comparisons with resting is indicated as follows: ***, $P \leq 0.001$; *, $P \leq 0.05$. Standard deviation represents biological replicates.

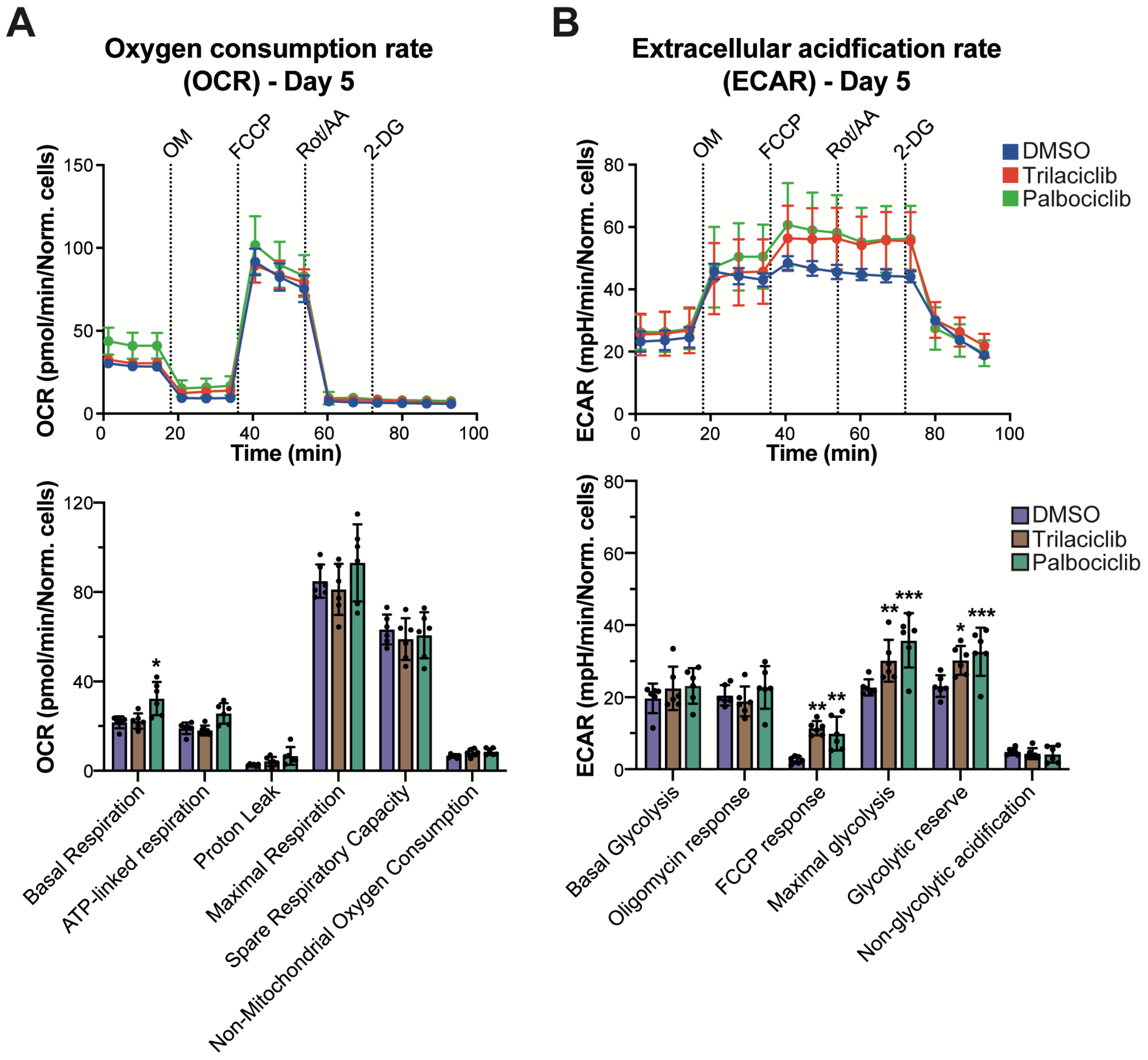


**Figure S7. Trilaciclib enhances the Warburg effect in A549 cells. A)** Oxygen consumption rate (OCR) and **B)** extracellular acidification rate (ECAR) in trilaciclib-, palbociclib- and DMSO-treated K562 cells after 5 days. The statistical significance of the comparisons with resting is indicated as follows: ***, P ≤ 0.001; **, P ≤ 0.01; *, P ≤ 0.05. Standard deviation represents biological replicates.

**R Script.**

```r
####  Load libraries  ####
library(data.table)
library(tidyr)
library(corrplot)
library(ggplot2)
library(ggrepel)
library(concaveman)
library(plyr)
library(dplyr)
library(factoextra)
library(ggfortify)
library(plotly)
library(rgl)
library(plotly)
library(rgl)
library(limma)
library(qvalue)
library(EnhancedVolcano)
library(ggrepel)
library('org.Hs.eg.db')

####  Load DIA-NN data  ####
pg<-fread('report.pg_matrix.tsv')
pr<-fread('report.pr_matrix.tsv')

####  Add number of peptides per protein entry  ####
t <- table(unique(pr[,c('Protein.Group','Stripped.Sequence')])$Protein.Group)
pg$Peptide.Count <- t[match(pg$Protein.Group,names(t))]
write.table(pg,"ProteinGroup.txt",sep = "\t",
        row.names = F,quote=F)
####  Read 'proteinGroup.txt' file  ####
df.prot = read.table("ProteinGroup.txt",header=T,sep="\t",stringsAsFactors = F,
              comment.char = "",quote ="")

####  Filter hits with at least 2 peptides  ####
df.prot = df.prot[!df.prot$Peptides<=1,]

####  Log2 transformation  ####
# Select columns of interest #
df.LFQ = df.prot[6:11]

####  Change NA by 0  ####
df.LFQ[df.LFQ==0] <- NA
```

```
####  Add a column with Identifier  ####
# Select columns of interest #
rownames(df.LFQ) = df.prot$Identifier
df.LFQ[1:6] <- log(df.LFQ[1:6], 2)

####  Filter two of three NA values in at least one group  ####
# Select columns of interest #
df.LFQ$na_count_DMSO = apply(df.LFQ,1,function(x) sum(is.na(x[1:3])))
df.LFQ$na_count_Trilaciclib = apply(df.LFQ,1,function(x) sum(is.na(x[4:6])))
df.LFQ.filter = df.LFQ[df.LFQ$na_count_DMSO<1|df.LFQ$na_count_Trilaciclib<1,1:6]

#### Read 'Metadata.txt' file ####
# two column file: 'Sample' and 'Group'
# 'Sample' column with same names than samples in 'ProteinGroup.txt'
# 'Group' column with group names, eg.: 'Control' and 'Trilaciclib'
metadata = read.table("Metadata.txt", stringsAsFactors = FALSE,
                   header = TRUE, sep = "\t")
conds = as.factor(metadata$Group)
scale_color_manual = c("#c9ac63","#62a18a")[as.factor(conds)]
names(scale_color_manual) = conds

#### Median normalization ###
filter <- df.LFQ.filter[,c(1,1:6)]
filter  [filter  == -Inf] <- NA
filter <- filter [,-1]
suppressMessages(suppressWarnings(library(limma, quietly = TRUE)))
filter_median_norm = normalizeMedianValues(filter)
filter_median_norm_plot <- gather(as.data.frame(filter_median_norm, sample, Intensity))
filter_median_norm_plot <- na.omit(filter_median_norm_plot)
filter_median_norm.df <- as.data.frame(filter_median_norm)
median_norm <-filter_median_norm.df
write.table(median_norm,"normalised.txt",sep = "\t",
          row.names = F,quote=F)

####  Intensities boxplot after normalisation  ####
boxplot(median_norm,
        col=scale_color_manual,
        las=2,
        main="Intensity Log2",
        cex.main=2,
        cex.axis=0.6)

####  Density plot after normalisation  ####
plotDensities(median_norm,
            main="Density (log2)",
            col=scale_color_manual,
            legend="topright")
```

```
#### Correlation plot after normalization ####
MultiScatter=cor(median_norm, use="pairwise.complete.obs")
corrplot(MultiScatter, type="upper",
        method = "color",
        col=colorRampPalette(c("lightskyblue","navy"))(100),
        is.corr=FALSE,
        tl.col="black",
        tl.cex=1,
        number.cex=0.6,
        number.digits=3,
        addgrid.col = "black"
)

#### Dendrogram ####
d <- dist(t(na.omit(median_norm))) ## distance between samples
hc <- hclust(d) ## hierarchical clustering
plot(hc) ## visualisation

#### PCA Plots ####
pca_res <- prcomp(t(na.omit(median_norm)), scale = TRUE)
fviz_eig(pca_res, addlabels = TRUE, linecolor = "black",
        barcolor = "black", barfill = "#1bbcf2")
conds = as.factor(metadata$Group)
cond_colours2 = c("black","red","blue")[as.factor(conds)]
names(cond_colours2) = conds
pca_data=prcomp(t(na.omit(median_norm)))
pca_data_perc=round(100*pca_data$sdev^2/sum(pca_data$sdev^2),1)
df_pca_data=data.frame(PC1=pca_data$x[,1], PC2=pca_data$x[,2], PC3=pca_data$x[,3],
sample=colnames(median_norm), condition = metadata$Group)
find_hull=function(df_pca_data) df_pca_data[chull(df_pca_data$PC1,df_pca_data$PC2),]
hulls=ddply(df_pca_data, "condition", find_hull)

# PC1 vs PC2
pca_data=prcomp(t(na.omit(median_norm)))
pca_data_perc=round(100*pca_data$sdev^2/sum(pca_data$sdev^2),1)
pca_data_perc=round(100*pca_res$sdev^2/sum(pca_res$sdev^2),1)
df_pca_data=data.frame(PC1=pca_data$x[,1], PC2=pca_data$x[,2],
sample=colnames(median_norm), condition = metadata$Group)
find_hull=function(df_pca_data) df_pca_data[chull(df_pca_data$PC1,df_pca_data$PC2),]
hulls=ddply(df_pca_data, "condition", find_hull)
ggplot(df_pca_data, aes(PC1,PC2, color=condition, fill=condition))+
  geom_point(size=3) +
  #scale_color_manual(values = 'blue", "black", "red", "yellow")) +
  theme_bw()+
  labs(title= "PC1 vs. PC2", x=paste0("PC1(",pca_data_perc[1],"%)"),y=paste0("PC2
(",pca_data_perc[2],"%)")) +
  geom_text_repel(aes(label=metadata$Sample), point.padding = 0.5) +
```

```
  ggforce::geom_mark_ellipse(aes(label=metadata$Group, fill = metadata$Group), alpha = 0.1, con.type = "straight", con.border = "none", show.legend = FALSE,)

# 3D PCA
Treatments <- factor(metadata$Group, levels=c("Control", "Trilaciclib"), labels=c("Control", "Trilaciclib"))
df_pca_data=data.frame(PC1=pca_data$x[,1], PC2=pca_data$x[,2], PC3=pca_data$x[,3], sample=colnames(median_norm), condition = metadata$Group)
find_hull=function(df_pca_data) df_pca_data[chull(df_pca_data$PC1,df_pca_data$PC2),]
hulls=ddply(df_pca_data, "condition", find_hull)
t <- list(
  family = "sans serif",
  size = 14,
  color = toRGB("grey50"))
fig <- plot_ly(df_pca_data, x = ~PC2, y = ~PC1, z = ~PC3, text = metadata$Sample, textposition = 'middle right', shape = FALSE, label.size = 5, color = Treatments, colors = c("Control"="#5cb57b","Trilaciclib"="#9c3353"))
fig <- fig %>% add_markers()
fig <- fig %>% add_text(textposition = "top right")
fig <- fig %>% layout(legend=list(title=list(text='<b> Treatment </b>')))
fig <- fig %>% layout(scene = list(xaxis = list(linecolor='black',mirror = T, title = paste0("PC2(",pca_data_perc[2],"%)")),
                       yaxis = list(linecolor='black',mirror = T,title = paste0("PC1 (",pca_data_perc[1],"%)")),
                       zaxis = list(linecolor='black',mirror = T,title = paste0("PC3 (",pca_data_perc[3],"%)"))))
fig

#### LIMMA analysis ####
design = model.matrix(~0+Group, data=metadata)
# fit linear model to the data
fit <- lmFit(median_norm, design)
# pairwise comparison between control vs treatment
cont <- makeContrasts(GroupTrilaciclib-GroupControl, levels = design)
fit2 = contrasts.fit(fit,contrasts = cont)
# this is the final results array- the standard errors are moderated using a simple empirical Bayes model using eBayes.
# The eBayes command will compute the consensus pooled variance, and then use it to compute the empirical Bayes (moderated) pooled variance for each gene.
# This also adjusts the degrees of freedom for the contrast t-tests.
# The command also computes the t-tests and associated p-values.
fit3 <- eBayes(fit2)
finalresult <- topTable(fit3, adjust.method = "BH", number = Inf)
setDT(finalresult, keep.rownames ="Identifier")
write.table(finalresult,"finalresult.txt",sep = "\t", row.names = F,quote=F)
setDT(df.LFQ.filter, keep.rownames ="Identifier")
write.table(df.LFQ.filter,"LFQ_filter.txt",sep = "\t",
          row.names = F,quote=F)
```

```
merged_finalresult <- merge(finalresult,df.LFQ.filter,by="Identifier") # merge with original dataset from MQ to have all additional info, eg number of peptides etc
merged_finalresult <- merge(merged_finalresult,df.prot,by="Identifier") # merge with original dataset from MQ to have all additional info, eg number of peptides etc
write.table(merged_finalresult,"mergedfinalresult.txt",sep = "\t",
        row.names = F,quote=F)
summary(decideTests(fit3))

# q-value
pvalue.df = merged_finalresult[,c(5)]
pvalue.df[pvalue.df==0] <- NA
rownames(df.prot) = pvalue.df$Identifier
pvalue.df <- qvalue(pvalue.df)
summary(pvalue.df)
hist(pvalue.df)
merged_finalresult$pvaluecheck <-pvalue.df$pvalues
merged_finalresult$qvalue <-pvalue.df$qvalues

# Volcano plot adj.q-value:
## add a column of NAs
merged_finalresult$diffexpressed <- "NO"
## if log2Foldchange > 0.59 and adj.P.Val < 0.05, set as "UP"
merged_finalresult$diffexpressed[merged_finalresult$logFC > 0.59 & merged_finalresult$qvalue < 0.05] <- "UP"
## if log2Foldchange < -0.59 and adj.P.Val < 0.05, set as "DOWN"
merged_finalresult$diffexpressed[merged_finalresult$logFC < -0.59 & merged_finalresult$qvalue < 0.05] <- "DOWN"

merged_finalresult$delabel <- NA
merged_finalresult$delabel[merged_finalresult$diffexpressed != "NO"] <- merged_finalresult$Genes[merged_finalresult$diffexpressed != "NO"]
## Plot the points with "diffexpressed"
volcano <- ggplot(data=merged_finalresult, aes(x=logFC, y=-log(qvalue), col=diffexpressed, label=delabel)) +
  geom_point() +
  ## geom_text()+
  geom_text_repel() +
  theme_classic() +
  xlab("Log2 fold change") +
  ylab("-Log10 adj p-value with q-value")
volcano <-volcano + geom_vline(xintercept = 0.59, linetype="dashed", col = "grey", size=0.5)
volcano <-volcano + geom_vline(xintercept = - 0.59, linetype="dashed", col = "grey", size=0.5)
volcano <-volcano + geom_hline(yintercept = -log(0.05), linetype="dashed", col = "grey", size=0.5)
volcano <-volcano + scale_color_manual(values = c("#709ec2", "black", "#f2a257"))
volcano

#### Gene Set Enrichment Analysis (GSEA)  ####
```

```r
# Set the desired organism
# CSV file containing a list of gene names and log2 fold change values.
# Use the `Genes´ for the index in the title, and `logFC´for the log2 fold change values.
# Sort the list of log2 fold change values in decreasing order.
# Download the The Molecular Signatures Database (MSigDB) from https://www.gsea-msigdb.org/gsea/msigdb/index.jsp
GSEA_input = read.table("GSEA_input.txt",header=T,sep="\t",stringsAsFactors = F,
                        comment.char = "",quote ="")
gmt = read.gmt("h.all.v7.0.symbols.gmt")
gmt$ont = gsub("HALLMARK_", "", gmt$ont)
genelist <- GSEA_input[,c(2,3)]
genelist1 = pull(genelist,logFC)
names(genelist1) = pull(genelist, Genes)
genelist1 = sort(genelist1, decreasing = TRUE)
gsea = GSEA(geneList=genelist1,
       exponent = 1,
       nPerm = 10000,
       minGSSize = 3,
       maxGSSize = 1000,
       pvalueCutoff = 0.05,
       pAdjustMethod = "BH",
       verbose = TRUE,
       TERM2GENE = gmt,
       by = "fgsea")
gsea_result = as_tibble(gsea)
gsea_result

write.table(gsea_result,"GSEA_result_Hallmark.txt",sep = "\t",
       row.names = F,quote=F)
# Use the `Gene Set` param for the index in the title, and as the value for geneSetId
dotplot(gsea, showCategory=1, split=".sign") + facet_grid(.~.sign)
```
